# Boundary conditions for early life converge to an organo-sulfur metabolism

**DOI:** 10.1101/487660

**Authors:** Joshua E. Goldford, Hyman Hartman, Robert Marsland, Daniel Segrè

## Abstract

It has been suggested that a deep memory of early life is hidden in the architecture of metabolic networks, whose reactions could have been catalyzed by small molecules or minerals prior to genetically encoded enzymes (1–6). A major challenge in unraveling these early steps is assessing the plausibility of a connected, thermodynamically consistent proto-metabolism under different geochemical conditions, which are still surrounded by high uncertainty. Here we combine network-based algorithms (9, 10) with physicochemical constraints on chemical reaction networks to systematically show how different combinations of parameters (temperature, pH, redox potential and availability of molecular precursors) could have affected the evolution of a proto-metabolism. Our analysis of possible trajectories indicates that a subset of boundary conditions converges to an organo-sulfur-based proto-metabolic network fueled by a thioester- and redox-driven variant of the reductive TCA cycle, capable of producing lipids and keto acids. Surprisingly, environmental sources of fixed nitrogen and low-potential electron donors seem not to be necessary for the earliest phases of biochemical evolution. We use one of these networks to build a steady-state dynamical metabolic model of a proto-cell, and find that different combinations of carbon sources and electron acceptors can support the continuous production of a minimal ancient “biomass” composed of putative early biopolymers and fatty acids.

---

The structure of metabolism carries a memory of its evolutionary history that may date back to before the onset of an RNA-based genetic system (1–6). Decoding this ancient evolutionary record could provide important insight into the early stages of life on our planet (2, 5–8), but constitutes a challenging problem. This challenge is due both to the difficulty of interrogating complex biochemical networks under different environmental conditions, and to the uncertainty about these conditions on prebiotic Earth. Estimates of plausible Archean environments that led to the emergence and evolution of living systems vary dramatically (21, 22), ranging from alkaline hydrothermal vents driven by chemical gradients (23) to acidic ocean seawater driven by photochemistry (3, 4). Although geochemical data support the availability of mid-potential electron donors (H_2_) (24), sulfur (H_2_S) and potentially fixed carbon (25) in ancient environments, several key molecules used in living systems may have been severely limiting, including a source of fixed nitrogen (26, 27) (e.g. ammonia), low-potential electron donors (28, 29) and phosphate (30–32). Rather than assuming a steady supply of these biomolecules, one can critically revisit the notion that these molecules would have been necessary for the emergence of ancient proto-metabolic systems. Indeed, we recently used network-based algorithms to ask whether early metabolism would have required a source of phosphate, and found evidence that thioesters, rather than phosphate, may have endowed ancient metabolism with key energetic and biosynthetic capacity (12). This raises the broader question of whether other molecules and physico-chemical conditions may not be as crucial as previously thought for the emergence of a proto-metabolism. Understanding these dependencies could also reveal whether specific components of metabolism are particularly robust or fragile with respect to these initial conditions.

A computational method that can help address these questions is the network expansion algorithm, which simulates the growth of a biochemical network by iteratively adding to an initial set of compounds the products of reactions enabled by available substrates, until convergence (9, 10). This algorithm, in its application to the study of ancient life (11)(12), relies on three key assumptions: *first*, that chemical reactions successful in early processes were gradually augmented with new pathways, but never replaced; *second*, that, over long time-scales, horizontal gene transfer produced abundant shuffling of biochemical reactions across different organisms (19, 20), suggesting that an ecosystem-level approach to metabolism may be particularly suitable for describing ancient biochemistry; and *third*, that inorganic or small molecular catalysts (12, 13) could catalyze, in a weaker and less specific manner relative to modern enzymes, a large number of metabolic reactions, as confirmed by an increasing body of experimental evidence (14–18).

In this paper, we systematically explore a combinatorial set of molecules and parameters associated with possible early Earth environments, and use an enhanced network expansion algorithm to determine which proto-metabolic networks are thermodynamically reachable under each of these initial conditions. We further use constraint-based flux balance modeling to demonstrate the capacity of some of these networks to sustain flux, in a way that resembles homeostatic growth of present-day cells. Our results suggest that a thioester-driven organic network may have robustly arisen without phosphate, fixed nitrogen or low-potential electron donors. This network, by supporting the biosynthesis of keto acids and fatty acids may have prompted the rise of complex self-sustaining biochemical pathways, marking a key transition towards the origin of life.

We first sought to systematically characterize the effect of various geochemical scenarios on the possible structure of ancient metabolism. Building on prior work (11, 12), we constructed a model of ancient biosphere-level metabolism based on the KEGG database (33). We first modified the network (as described in Methods) to account for previously proposed primitive thioester-coupling and redox reactions (12). For each possible set of environmental parameters (including temperature, pH and redox potential) we computed the thermodynamic feasibility of each reaction, and removed infeasible reactions (see Methods). This allowed us to implement a thermodynamically-constrained network expansion algorithm (12), which iteratively adds metabolites and thermodynamically-feasible reactions to a network until convergence (see Methods). We performed thermodynamically-constrained network expansion (see Methods and Fig. 1A) for *n*=672 different geochemical scenarios, systematically varying pH, temperature, redox potential of primitive redox systems, and the availability of key biomolecules including thiols (that subsequently form thioesters), fixed carbon (formate/acetate) and fixed nitrogen (ammonia) (Methods, Fig. 1).

**Figure 1:**
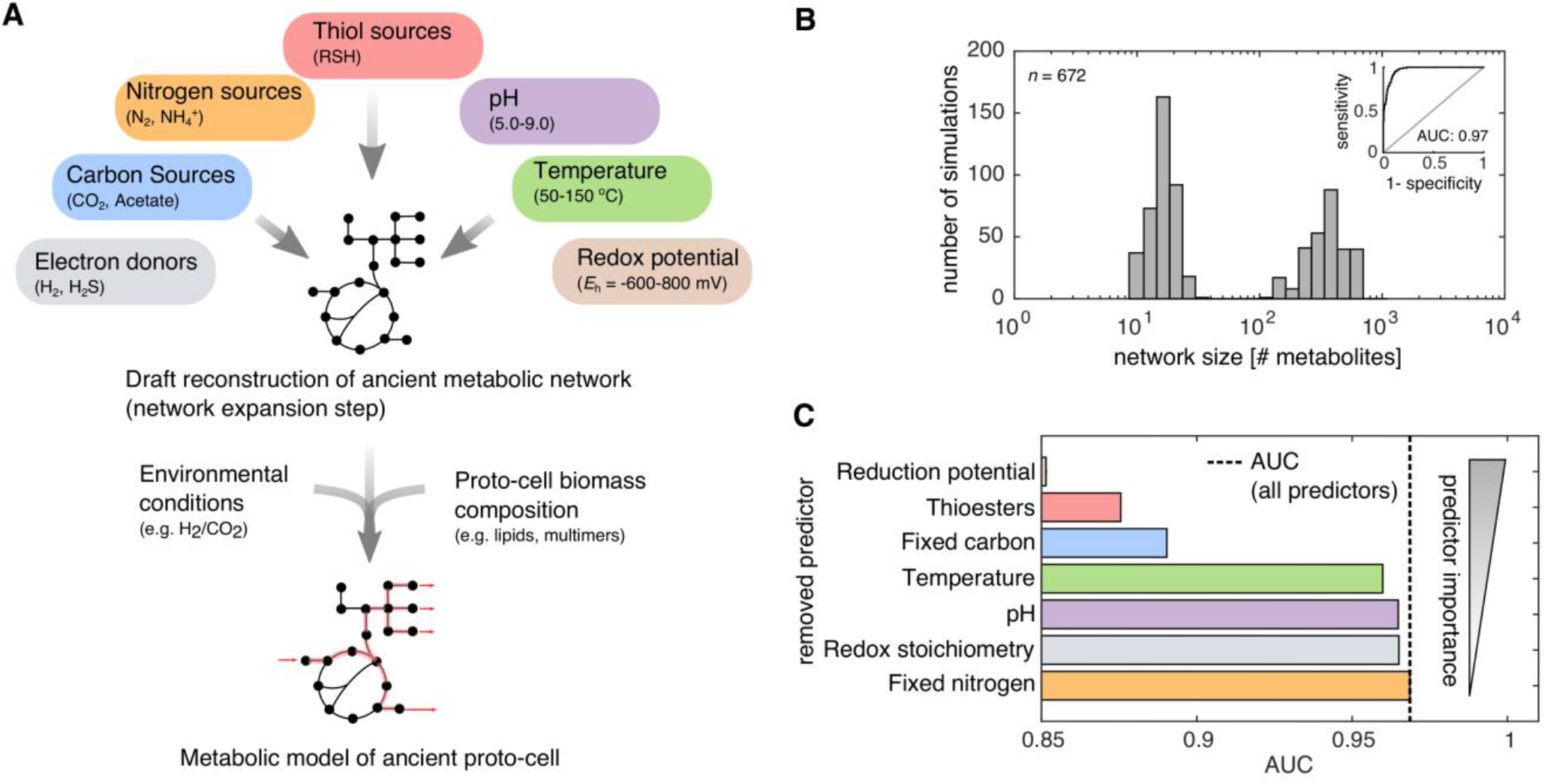
Nitrogen is not essential for the initial expansion of metabolism. (A) A thermodynamically-constrained network expansion algorithm was used to simulate the early expansion of metabolism under 672 scenarios, systematically varying the availability of reductants in the environment, pH, carbon sources, the presence of thiols, temperature and the availability of ammonia. (B) A histogram of network sizes (*x*-axis, number of metabolites) revealed that 43 % (288/672) of the scenarios resulted a bimodal distribution, where expansion occurred beyond 100 metabolites. (inset) A logistic regression classifier was constructed to predict whether a geochemical scenario resulted in a network that exceeded 100 metabolites, and a receiver operating curve (ROC) was plotted. The trained classifier resulted in an area under the curve (AUC) of 0.97 and leave-one out cross-validation accuracy of 0.89. (C) Models were trained without information on specific geochemical variables (*y*-axis), and the ensuing AUC was plotted as a bar-chart (*x*-axis), revealing that knowledge of the availability of fixed nitrogen offers no information on whether networks expanded.

Of the 672 different simulated geochemical scenarios, we found that 288 (43%) expanded to networks containing over 100 metabolites (Fig. 1B). A logistic regression classifier that uses geochemical parameters as predictors (see Methods and Fig. 1C) allowed us to quantify the importance of each environmental parameter in determining whether the expanded network would reach such a large size. Surprisingly, removing the variable associated with presence/absence of ammonia did not affect predictive power of the classifier, suggesting that a source of fixed nitrogen is not an important determinant of the expansion. Consistent with the relevance of this result to ancient metabolism, we found that the enzymes that catalyze reactions in the expanded networks before the addition of ammonia were depleted in nitrogen-containing coenzymes (see Fig. S1C-D, one-tailed Wilcoxon sign rank test: *P* < 10^−24^) and in active site amino acids with nitrogeneous side chains (see Fig. SE-F, one-tailed Wilcoxon sign rank test: *P* < 10^−24^) relative to enzymes added after the addition of ammonia (see Supplemental Text). These results suggest that ammonia may have not been essential for the initial expansion of metabolism, and point to a thioester-coupled organo-sulfur metabolic network (Fig. 1) as a core network that deserves further attention.

Beyond the dispensability of nitrogen, the simulations described above revealed a number of relationships between plausible geochemical scenarios and the structure and size of our simulated proto-metabolic networks. First, expansion beyond 100 metabolites was feasible in the absence of a source of fixed carbon, but only when thiols were provided in the seed set, highlighting the importance for thioester-coupling for ancient carbon fixation pathways (12, 28, 29). The presence of thiols enabled the production of key biomolecules, including fatty acids and branched chain keto acids (see Fig S2). Second, we explored the effect of the primitive redox system by systematically varying the reduction potential of the electron donor in the seed set (see Methods, Fig. 2A). Unexpectedly, we found that as we increased the fixed potential of the electron donor, expansion to a large network was feasible over a broad range of reduction potentials (between −150 and 50 mV). Only upon reaching 50mV the expanded network collapsed to a much smaller solution, suggesting that the generation of low-potential electron donors from H_2_ may not have been a necessary condition for the early expansion of a proto-metabolism (Fig. 3A). Thus, less stringent constraints, e.g. the presence of mid-potential redox couples and thioester-forming thiols could have enabled the emergence of an autotrophic proto-metabolic network capable of producing key biomolecules. Notably, both functions can be potentially carried out by disulfides.

**Figure 2:**
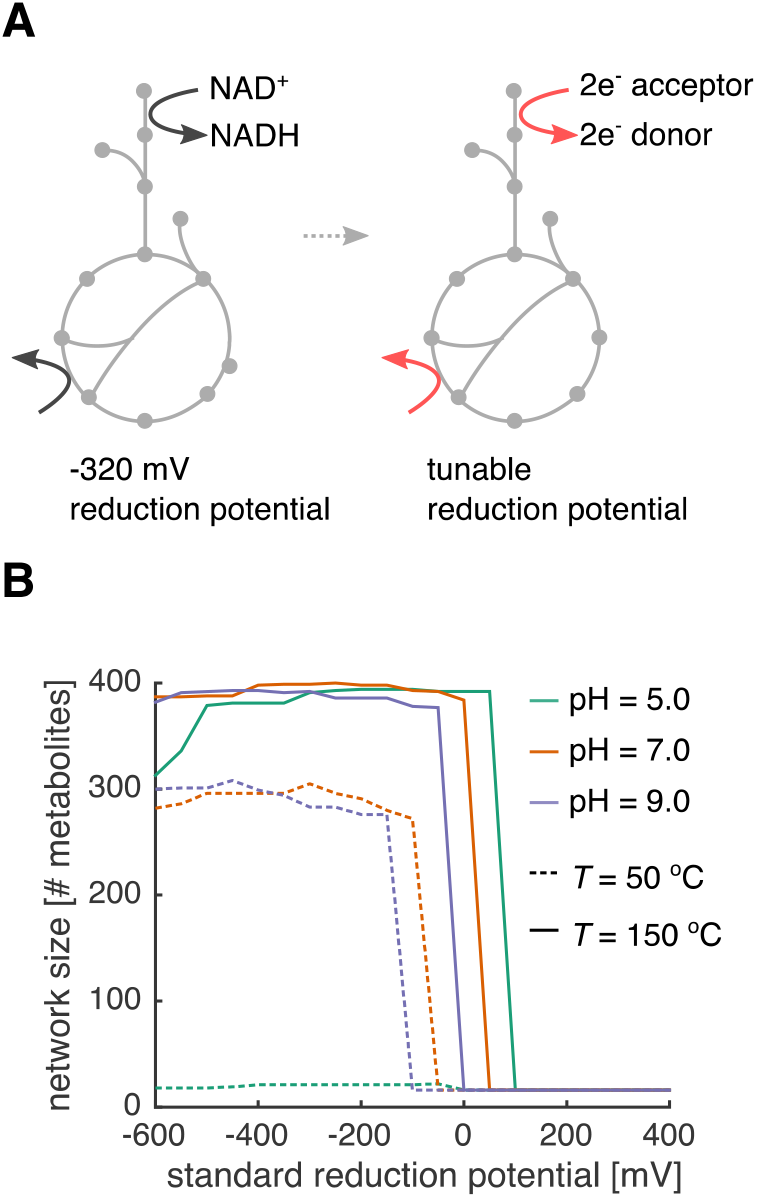
Reduction potential of nicotinamide and flavin substitutes influences network expansion. (A) Redox coenzymes (NAD, NADP, and FAD) were substituted with an arbitrary electron donor/acceptor at a fixed reduction potential. (B) We performed thermodynamic network expansion in acidic (pH 5), neutral (pH 7) and alkaline (pH 9) conditions at two temperatures (*T* =50 and 150 °C), using a two-electron redox couple at a fixed potential (*x*-axis) as a substitute for NAD(P)/FAD coupling in extant metabolic reactions (see Methods). We plotted the final network size across all pH and temperatures with no fixed carbon sources (e.g. only CO_2_) and thiols. Notably, for these simulations, we used a base seed set of: H_2_, H_2_S, H_2_O, HCO_3_^−^, H^+^ and CO_2_.

Analysis of the expanded networks without nitrogen revealed that a large number of different initial conditions converged to similar expanded organo-sulfur proto-metabolic networks, spanning variants of key pathways in central carbon metabolism (Fig. 3B). For the majority of simulations, variants of modern heterotrophic carbon assimilation pathways, including the glyoxylate cycle and TCA cycle, were highly represented in the network (Fig. 3B). Several carbon fixation pathways were highly represented in the simulated networks as well: in over half of the networks that expanded beyond 100 metabolites, we found 92 % (12/13) of the compounds (or generalized derivatives) that participate in the reductive tricarboxylic acid (rTCA) cycle, with the exception of phosphoenolpyruvate. We also found that under several geochemical conditions, all intermediates were producible for three carbon fixation pathways, including the 3-hydroxypropionate bi-cycle, the hydroxypropionate-hydroxybutylate cycle, and the dicarboxylate-hydroxybutyrate cycle (Fig. 3A). At-most, only 3 of 9 metabolites used in the Wood-Ljungdahl (WL) pathway were observed, due to the lack of nitrogen-containing pterins in the network. This does not necessarily rule out the primordial importance of the WL-pathway, as its early variants could have been radically different than today’s WL-pathway, relying on native metals to facilitate reduction of CO_2_ to acetate (25, 29). In addition to observing a large number of metabolites used in carbon fixation pathways, we found that a large fraction of the β-oxidation pathway was represented in our networks, which may have supported the production of fatty acids in ancient living systems by operating in the reverse direction. Lastly, we observed that the majority of intermediates involved in the production of branched-chain amino acids were also producible in the expanded networks.

**Figure 3:**
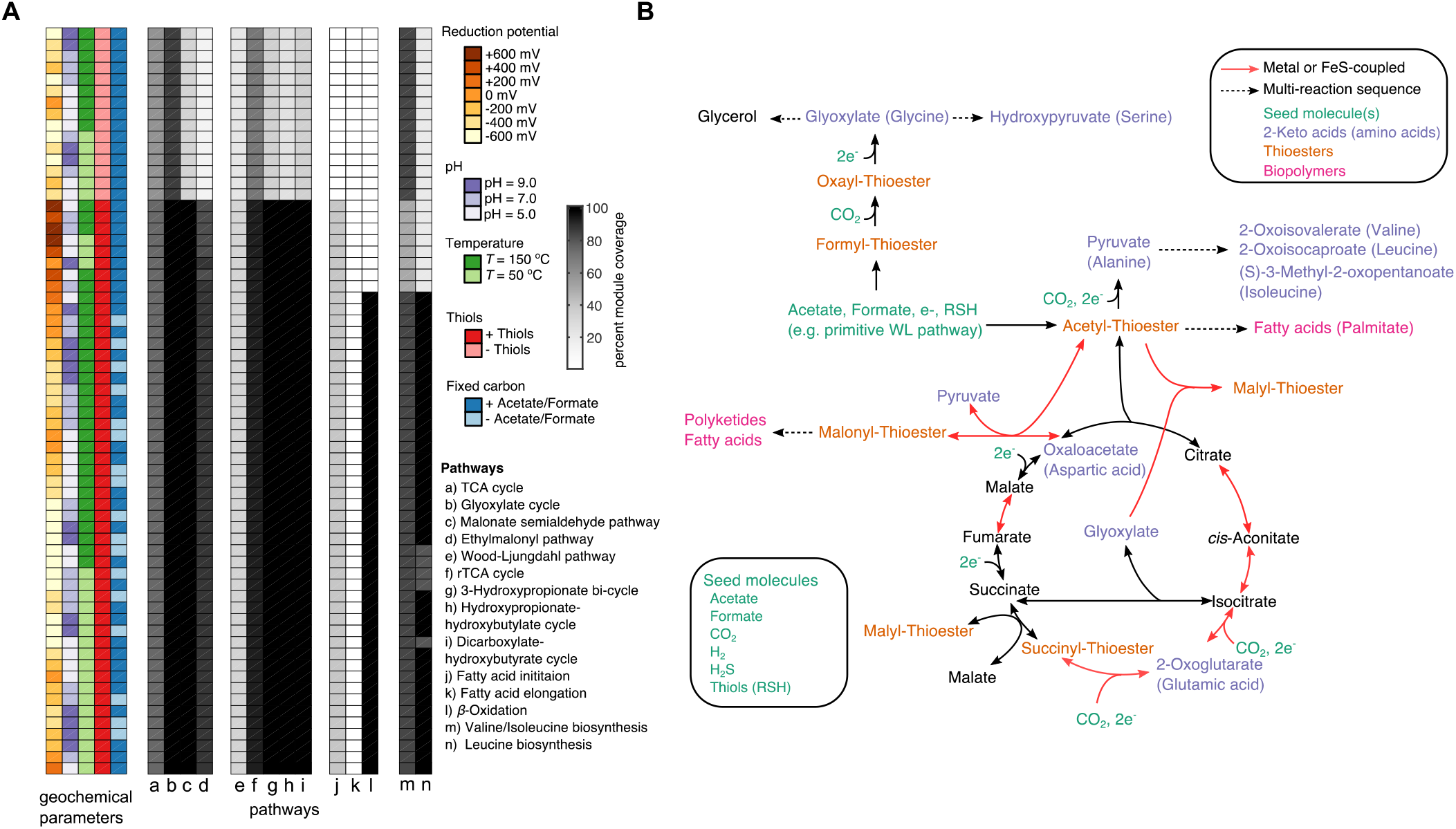
Systematic exploration of prebiotic scenarios reveals a core organo-sulfur network. (A) A thermodynamically constrained network expansion algorithm was used to simulate the early expansion of proto-metabolism under various scenarios, including the availability of reductants in the environment, pH, temperature, and the availability of fixed carbon sources and thiols. The proportion of molecules selected KEGG modules involved carbon metabolism are plotted as a heatmap to the right of the parameters. (B) A representation of the core network producible from a prebiotically plausible seed set without nitrogen or phosphate (bottom left box). Acetyl-thioesters are first produced, potentially from a primitive Wood-Ljungdahl pathway (8, 25) from acetate and thiols provided as seed molecules (green). Acetyl-thioesters enable the production of all intermediates in the reductive tricarboxylic acid (rTCA) cycle, with the exception of phosphoenolpyruvate. ATP-dependent reactions in the rTCA cycle may have been substituted with a primitive malate sythase and transthioesterification of succinate as well as the recently discovered reversible citrate synthase (45, 46). The keto acid precursors for 8 common amino acids (A,D,E,G,I,L,S,V) are highlighted in purple, while routes to thioester-mediated polymerization of fatty acids and polyketides are highlighted in pink.

A more detailed analysis of the convergent organo-sulfur proto-metabolic network reveals new possible ancestral metabolic pathways that involve previously unexplored combinations of reactions and metabolites. Fig. 3B shows a variant of the (r)TCA cycle that is a component of these expanded networks, and that may have served as the core organo-sulfur network fueling ancient living systems. Rather than using ATP-dependent reactions found in extant species (e.g. Succinyl-CoA synthetase and ATP citrate lyase), these reactions are substituted with non-ATP-dependent reaction mechanisms. For instance, the production of a succinyl-thioester in the extant rTCA cycle relies on Succinyl-CoA synthetase, performing the following reaction: ATP + Succinate + CoA ➔ Succinyl-CoA + ADP + P_i_. However, in the network presented in Fig. 3B, malyl-thioester, producible through alternative reactions, donates a thiol to succinate, subsequently forming a succinyl-thioester. This (r)TCA cycle analogue is able to produce eight keto acids normally serving as key intermediates and precursors to common amino acids in central carbon metabolism (glyoxylate, pyruvate, oxaloacetate, 2-oxoglutarate and hydroxypyruvate), as well as a few branched-chain keto acids. Additionally, long-chain fatty acids like palmitate are producible in this network, driven by thioester and redox-coupling rather than ATP, like in extant fatty acid biosynthesis. Thus, despite the simplicity of seed compounds, several small molecular weight keto acids and fatty acids may have been producible in an organo-sulfur proto-metabolism.

So far, we have focused only on the topology and global thermodynamic feasibility of putative ancient metabolic networks. Inspired by recent studies on the molecular budget of present-day cells, we decided to further explore whether proto-metabolic networks could support steady state fluxes, and fuel primitive proto-cells with internal energy sources (e.g. thioesters), redox gradients, and primitive biopolymer capable of catalysis and compartmentalization. Flux balance analysis (FBA), originally developed for the study of microbial metabolism, enables the prediction of systems-level properties of metabolic networks at steady-state (34). Fundamentally, FBA computes possible reaction rates in a network constrained by mass and energy balance, usually under the assumption that a specific composition of biomolecules is efficiently produced during a homeostatic growth process. In microbial metabolism, FBA is used to simulate the production of cellular biomass (e.g. protein, lipids, and nucleic acids) at fixed proportions, which are derived from known composition of extant cells. We realized that the same approach could help test the sustainability of a proto-metabolic biochemical system, provided that we could develop a plausible hypothesis for the “biomass composition” of ancient proto-cells. As a starting point, we recalled Christian de Duve’s suggestion that the thioester-driven polymerization of monomers producible from ancient proto-metabolism may have led to “catalytic multimers”, which could have served as catalysts for ancient biochemical reactions (4). Under nitrogen limited conditions, keto acids producible from proto-metabolism (see Fig. 3B) could have been reduced to α-hydroxy acids, and polymerized into polyesters using thioesters as a condensing agent (see Fig. S3). Recent work has suggested that polymers of α-hydroxy acids may have been stably produced in geochemical environments (35), and that these molecules could have served as primitive catalysts (36). These results all point to the intriguing possibility that the thioester-driven polymerization of α-hydroxy acids (producible from keto acid precursors of common amino acids) generated the first metabolically sustainable cache of ancient catalysts, leading to a collectively autocatalytic protocellular system. We employed FBA to specifically test the feasibility of such a system. Using an expanded metabolic network as a scaffold for network reconstruction (Fig. 3B), we constructed a constraint-based model of an ancient proto-cell using a biomass composition consisting of fatty acids (for proto-cellular membranes), “catalytic multimers” derived from eight keto acids (Fig. 4B), and redox and thioester-based free energy sources (Methods, Fig. 4A). We used thermodynamic metabolic flux analysis (TMFA), a variant of FBA that explicitly considers thermodynamic constraints (37) (see Methods), to determine whether homeostatic growth of the whole system was achievable. We found that growth of the proto-cell metabolic model is indeed feasible under a wide variety of assumptions regarding macromolecular compositions and input molecules (Fig. 4B). Notably, growth is achievable in simple chemoautotrophic conditions with either H_2_ or H_2_S, but not Fe(II), as electron donors (Fig. 4B). In this model, thiols and thioesters are not supplied as food sources, but rather are recycled during steady-state growth of the proto-cell. This reflects the possibility that thiols could have been initially supplied abiotically, followed by the rapid takeover of biotic production of mercaptopyruvate, a keto acid that could have been incorporated into primitive multimers.

**Figure 4:**
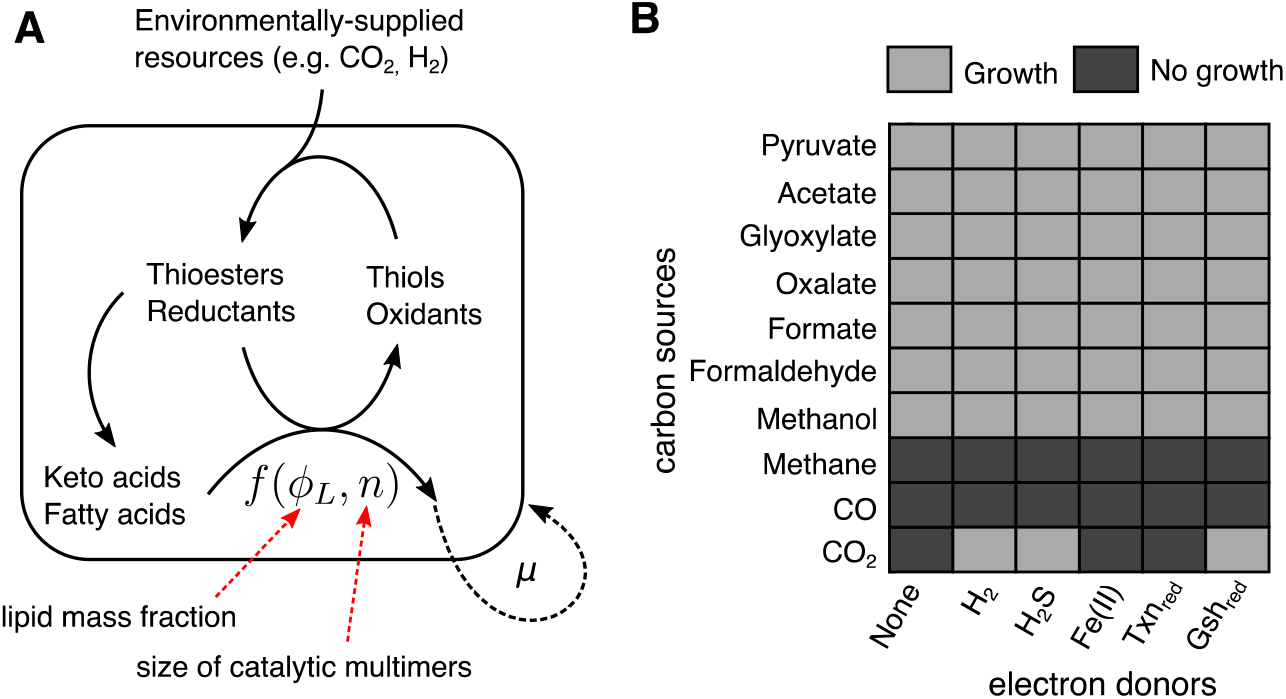
Constraint-based modeling of plausible ancient proto-cells. (A) We constructed a metabolic of model of plausible ancient proto-cell and used thermodynamic metabolic flux analysis (37) to simulate the maximum growth yields, steady-state fluxes, and metabolite concentrations under a variety of environmental conditions. The metabolic model was constructed using internally-generated reductants and thioesters that fueled biomass formation. The biomass composition was specified as variable fractions of fatty acids, and polymerized hydroxyacids from keto-acid precursors (see Methods). In this model, the internal redox coenzyme was assumed to be disulfide/dithiol at a standard reduction potential of -220 mV, and the production of biomass was fueled by the hydrolysis of acetyl-thioesters. We parameterized the biomass composition using a two-parameter model (see Methods), with the mass fraction of lipids in the proto-cell set to *ϕ_L_* = 0.2, and the the average size of a catalytic multimer, *n* = 10. (B) We simulated growth on a variety of simple carbon sources (*y*-axis) and electron donors (*x*-axis), and show environments supporting non-zero growth (light grey) or no growth (dark grey). Interestingly, H_2_, H_2_S and glutathione were the only reductants capable of supporting fully autotrophic growth on CO_2_. Furthermore, CO and Methane could not support growth in this model, while other one-carbon sources like methanol, formate and formaldehyde could support biomass growth.

While most efforts to reconstruct ancient biochemistry have traditionally relied on building qualitative models of small pathways (2, 4, 5, 7, 38), we found that quantitative modeling of larger networks can provide substantial new insight into the origin of life. By computationally mapping geochemical scenarios to plausible ancient proto-metabolic structures, we estimated which portions of extant biochemistry may have been very sensitive or very robust to initial geochemical conditions. Our approach reveals that, contrary to expectations (8, 28, 39), environmental sources of fixed nitrogen and low-potential electron donors may have not been necessary for early biochemical evolution, and a substantial degree of complexity may have emerged prior to incorporation of nitrogen into the biosphere (3). The key catalytic role played by nitrogen in the active sites of modern enzymes may have been preceded by positively charged surfaces or metal ions (18, 25), which could have been replaced by amino/keto acids with nitrogen side chains once nitrogen became incorporated into proto-metabolism. Our simulations also cast doubts on the essential role of a low-potential electron donor in early life (8, 28, 29), consistent with the proposal that low-potential electron donors may not be necessary for acetogenesis (40), and with the possibility that energy conservation via electron bifurcation might not have been necessary in primordial metabolism. The independence of our inferred ancestral networks of low-potential electron donors and ATP, both key substrates for nitrogen fixation (41), suggests that nitrogen fixation may have evolved later throughout the history of life (42–44). A striking feature of our analysis is the convergence of multiple geochemical scenarios towards a core organo-sulfur proto-metabolic network capable of producing various keto acids and fatty acids (Fig. 3B). This feature provides a window into how thioester-driven polymerization of α-hydroxy acid monomers (derived from producible keto acids) could have added primitive macromolecular organic catalysts (4) to initial inorganic minerals or metal ion catalysts (18, 25). Further tests of this hypothesis could be pursued by measuring the capacity of these polymers to catalyze key reactions in the network, and by exploring whether these organic compounds are produced in living systems today via mechanisms similar to polyketide or non-ribosomal peptide synthesis. Finally, our constraint-based models of this core organo-sulfur proto-metabolism provide a first example of how network expansion-based predictions can be translated into dynamical models, whose capacity to estimate sustainable collective growth can drive the search for specific self-reproducing chemical networks and metabolically-driven artificial protocells.

## Acknowledgements

We thank all members of the Segre Lab for helpful discussions. We acknowledge support by the Directorates for Biological Sciences (BIO) and Geosciences (GEO) at the NSF and NASA under Agreements No. 80NSSC17K0295, 80NSSC17K0296 and 1724150 issued through the Astrobiology Program of the Science Mission Directorate.

## Contributions

J.E.G., H.H. and D.S. designed the research. J.E.G. wrote code, ran simulations and performed analysis. R.M. contributed to the non-equilibrium steady-state modeling. J.G. and D.S. wrote the manuscript. All authors read and approved the final manuscript.

