## Supplementary material for "Boundary conditions for early life converge to an organo-sulfur metabolism"

#### This PDF file includes:

Supplementary text
Figs. S1 to S4
Caption for Database S1
References for SI reference citations

#### Other supplementary materials for this manuscript include the following:

Database S1

### Materials and methods

**Reconstruction of biosphere-level metabolic network.** Biosphere-level metabolism was reconstructed from the KEGG database (1) according to protocol described previously (2). We modified the network in several ways to model primitive thioester-based metabolic network without nitrogen or phosphate. First, to simulate the availability of thiols capable of forming thioesters, we included Coenzyme A, Acyl-Carrier Protein and Glutathione into the seed set. However, to enforce the constraint that these metabolites could only be used in reactions as coenzymes (and not products or substrates), we prevented the degradation by removing KEGG reactions R10747, R02973 and R02972.

We next assigned standard molar free energies to reactions using eQuilibriator at a predefined pH (3). Next we substituted NAD, NADP and FAD-coupled reactions with an arbitrary redox couple. For example, if the redox reaction $X_{ox}$ + NADH $\rightarrow$ $X_{red}$ + NAD+ was swapped with electron donor with a redox potential of $E_0^+$ mV, we would use the following formula to adjust the standard molar free energy for the new reaction $r'$:

$$\Delta_{r'} G'^{\circ} = \Delta_r G'^{\circ} + nF(E_0^+ - E_0)$$

where $n$ are the number of electrons transferred in reaction $r$ and $F = 96.485$ kJ/V. Note that if we assumed that the electron donor/acceptor substitute was a two electron donor/acceptor, we did not change the stoichiometry in the reaction equation. However, in the case where the electron donor/acceptor substitute was a single electron donor/acceptor, we change the stoichiometric coefficients to $s_{cj} = 2$ for all reactions $j$, where $c$ represents metabolites NAD(H), NADP(H) and FAD(H2). For this work, we systematically varied the reduction potential $E_0^+$ and stoichiometry of the primitive redox coenzyme.

**Thermodynamically-constrained network expansion.** We performed network expansion using thermodynamic constraints in a different way than performed previously (2) Previously, we removed reactions above a predefined free energy threshold of $\tau = 30$kJ/mol (2). For this work, we computed the lowest reaction free energy possible using estimates for upper and lower bounds on metabolite concentrations, $u_i$ and $l_i$, and removed reactions with a positive reaction free energy. For a given biochemical reaction at fixed temperature and pressure, $\Delta_r G'$ is defined as:

$$\Delta_r G' = \Delta_r G'^{\circ} + RT\ln \prod_i a_i^{s_{ir}}$$

where the $\Delta_r G'^{\circ}$ is the free energy change of the reaction at standard molar conditions, $R$ is the ideal gas constant, $T$ is temperature, $a_i$ is the activity of metabolite $i$ and $s_{ir}$ is the stoichiometric coefficient for metabolite $i$ in reaction $r$. We fixed $a_i$ for each reactions according to the following rules:

$$a_i = \begin{cases} u_i, & \text{if } s_{ir} < 0, \\ l_i, & \text{if } s_{ir} > 0. \end{cases} \quad [1]$$

We then removed reactions with a $\Delta_r G' \geq 0$. For all simulations we assumed that $u_i = 10^{-1}$ M and $l_i = 10^{-6}$ M. Note that because we model each reaction independently, metabolite concentrations could be inconsistent . For instance, if metabolite $i$ is the substrate for reaction $a$ and a product for reaction $b$, then $x_i = u_i$ for reaction $a$ and $x_i = l_i$ for reaction $b$.

Using this procedure to systematically remove reactions that were considered to be thermodynamically infeasible, we performed network expansion (4–6) using the computational procedure described in (2).

**Parameters for network expansion.** We systematically studied the size and composition of networks under precise environmental conditions by varying (a) the reduction potential from the environment, (b) pH, (c) temperature, (d) the presence or absence thiols, (e) the inclusion of fixed carbon into the seed set and (f) the inclusion of fixed nitrogen into the seed set. We now discuss each of these parameters in more detail:

- *Reduction potential and stoichiometry.* A wide range of environmental conditions could have provided electron donors at various potentials: high potential redox pairs, with strong oxidants like Fe(III), may have been present in oceans at high concentrations, while strong reductants like $H_2$, disulfides, proto-ferredoxin, or reductive carboxylation of thioesters have been produced via serpentinization or geochemical analogues of primitive metabolic pathways (7). We substituted reactions coupled to NAD, NADP and FAD with a generic single or double electron donor and acceptor pair at a fixed potential. To prevent unbalanced electron transfer, we removed the following transhydrogenase reactions: R10159, R01195, R00112, R09520, R09748, R05705, R05706, R09662, R09750. We then created a single or double electron donor/acceptor pair with a fixed reduction potential, $E_0^+$, ranging from -600 to 600 mV.

- *pH* We modified the pH by setting reaction free energies at various pH's (5.0-9.0) using eQuilibriator (3) which relies on the component contribution method (8).

- *Temperature.* Temperatures were assumed to have been within a range of 50-150 °C, spanning estimates of ocean seawater temperature in the Archean (9), up to some alkaline hydrothermal vent systems (10).

- *Thiols.* In our model we provided thiols that serve as substitutes for coenzymes that form thioester bonds in extant metabolic networks. To this end, we provided Coenzyme A, acyl-carrier protein and Glutathione in the seed set, but removed key degradation reactions to ensure these compounds only served as coenzymes, rather than material sources, during network expansion (2).

- *Fixed nitrogen.* To study the consequences of adding or removing a source of fixed nitrogen as a seed compounds for network expansion, we either added or removed ammonia from the seed set prior to expansion.

In addition to parameters we varied, we kept constant two additional parameters that could be studied in future work:

- *Metabolite concentrations.* Metabolite concentrations were assumed to be within 1 $\mu$M - 100 mM. The upper bound estimate is consistent with recent experimental data showing that key metabolites (formate, methanol, acetate and pyruvate) can be produced near 100 mM (11). Although we do not have empirical evidence to suggest a reasonable lower bound on metabolite concentrations in ancient metabolic networks, we assumed that 1 $\mu$M, the estimated lower bound in today's cells (12), was also the lower bound in our model of ancient metabolism.

- *Reactions with no free energy estimate.* 53 % of the biosphere-level metabolic network reactions have no free energy estimate (4851 of 9074). For all simulations presented in this paper, we assumed these reactions were blocked and did not include them in the network.

**Generalized linear modeling of network expansion results.** To access the effects of various parameters on the outcome of network expansion, we used generalized linear models to construct logistic regression classifiers to predict whether or not the network expanded beyond 100 metabolites using a combination of predictors, including categorical variables encoding whether or not ammonia, thiols or fixed carbon was provided in the seed set, and continuous variables encoding the reduction potential, pH and temperature used in each simulation. We first define the response variable for simulation $i$ as $y_i$ where $y_i = 1$ if the simulation resulted in a network that expanded beyond 100 metabolites, and zero otherwise. For the set of simulations performed in Fig. 1 in the main text, we constructed a design matrix consisting of categorical variables representing the following scenarios:

1. $x_{N,i} \in \{0,1\}$: 1 if ammonia was included in the seed set, and 0 otherwise.

2. $x_{S,i} \in \{0,1\}$: 1 if thiols were included in the seed set, and 0 otherwise.

3. $x_{C,i} \in \{0,1\}$: 1 if fixed carbon (acetate/formate) was included in the seed set, and 0 otherwise.

4. $x_{H,i} \in \mathbb{R}_{>0}$: A continuous variable representing the pH. Note for our simulations, we only explored acidic (pH=5), neutral (pH=7) and alkaline (pH=9) regimes.

5. $x_{E,i} \in \mathbb{R}$: A continuous variable representing the reduction potential at standard molar conditions (a the specified pH listed above). For our simulations, we explored a wide range of standard molar reduction potentials (from -600 mV to +600 mV).

6. $x_{T,i} \in \mathbb{R}$: A continuous variable representing the temperature. For our simulations, we explored two temperatures: a high temperature regime ($T = 150$ °C), and a low temperature regime ($T = 50$ °C).

We next constructed the following generalized linear model to model whether the network expanded beyond metabolites:

$$\text{logit}(y_i) = \beta_0 + \beta_N x_{N,i} + \beta_S x_{S,i} + \beta_C x_{C,i} + \beta_H x_{H,i} + \beta_E x_{E,i} + \beta_T x_{T,i} \quad [2]$$

We fit the parameters ($\beta$) using the *fitglm.m* function in MATLAB 2015a, and a receiver operating curve (ROC) was generated using the *perfcurve.m* function. For results presented in Fig. 2C in the main text, individual predictors were removed in the generalized linear model presented above. To access whether the trained logistic model served as an accurate classifier, we performed leave-one out cross-validation by removing individual samples from the training set an testing the accuracy of the trained classifier on the removed sample. This procedure resulted in an cross-validation accuracy of 0.89.

**Constraint-based modeling.** We constructed a model of a autocatalytic network at steady state using a variant of constraint-based modeling of cellular metabolism called thermodynamic-based metabolic flux analysis (TMFA) (13). TMFA transforms the non-linear constraints induced by imposing thermodynamic consistency into mixed-integer linear constraints. In this section, we first describe (a) the construction of primitive biomass composition for a model of an ancient proto-cell and (b) the formulation of TMFA used in this analysis.

***Prebiotic biomass equation.*** We constructed a simple model for the macromolecular composition of primitive proto-cells, using empirical knowledge of extant cellular life. Since our metabolic model of proto-metabolism does not included macromolecular production of nucleotides (and thus a nucleic acid based genetic system), we assume that the primary role of proto-cellular metabolism was to initially produce components for a cellular membrane and catalysts. Building off of Christian de Duve's multimer hypothesis (14), we first propose that the biomass can be constructed using a simple two parameter model consisting of the mass fraction of lipids $\phi_L$ and the average length of each catalytic multimer $n$.

- *Lipid mass fraction.* The lipid content in modern cells is roughly 10% of the total dry mass (Bionumbers ID: 111209) (15), primarily composed of the fatty acid palmitate. For our analysis, we assume that palmitate represents the sole component of lipids. Future models could incorporate glycerol, which enables the production of glycerolipids. While phosphate is used in cellular membranes as a polar head group to produce amphiphilic molecules, primitive processes may have conjugated negatively charged organic acids (e.g. oxalate) to glycerol via a thioester-mediated synthesis mechanism to create ampiphilic lipid molecules resembling modern phospholipids. For our initial model, we simply propose that palmitate was the initial amphiphilic component of primitive membranes, where the negatively charged polar carboxylate ion was sufficient for forming a membrane, and assumed that proto-cells consisted of a lipid mass fraction of $\phi_L$.

- *Catalytic multimer mass fraction.* We propose that ancient catalysts were composed of inorganic molecules (e.g. iron-sulfur clusters, metal ions, mineral surfaces) chelated with multimers of $\alpha$-hydroxy-acids (see Fig. 4A in main text). For our model, we assume that the eight keto acid precursors producible from our network were the dominate monomers of ancient multimeric catalysts. We assume that the total mass fraction of these catalysts are $1 - \phi_L = \phi_C = \sum_k \phi_k$, where $\phi_k$ is the mass fraction of polymerized monomer $k$. For our analysis, we assumed that each monomers is uniformly distributed, so that $\phi_k$ = constant for all $k$. Additionally, since each monomer must be reduced to $\alpha$-hydroxy acids, there is linear relationship between the electron demand, $s_e$, and the number of molecules of monomers produced. The stoichiometric equivalents of electron donors are thus:

  $$s_e = 2\sum_k \frac{\phi_k}{M_k}$$

  where $M_k$ is the molar mass of monomer $k$.

- *Average size of catalytic multimers.* The average size of mulimeric catalysts sets the number of thioester bonds required for synthesis of catalytic multimers. For each polymer of size $n$, there are $n-1$ thioester bonds required. In our model, the total number of monomers are fixed to be: $\sum_k \frac{\phi_k}{M_k}$, where $M_k$ is the molecular weight for monomer $k$. Thus for a fixed monomer length $n$, we can compute the number polymers using the following formula:

  $$P(n) = \frac{1}{n}\sum_k \frac{\phi_k}{M_k}$$

  The thioester demand is thus $s_t(n) = (n-1)P(n)$, or:

  $$s_t(n) = \frac{n-1}{n}\sum_k \frac{\phi_k}{M_k}$$

  For our analysis we assumed a fixed polymer length of size $n = 10$ monomers.

Using these two parameters, we constructed the biomass equation for the proto-cellular model.

***Thermodynamic Metabolic Flux Analysis (TMFA).*** To simulate a thermodynamically-feasible steady-state behavior of this metabolic network, we used thermodynamic metabolic flux analysis (TMFA) (13). Briefly, TMFA transforms the non-linear constraints induced by imposing thermodynamic consistency into mixed-integer linear constraints. We first converted the model into an irreversible model by modeling each reaction as both forward and backward half reactions. We then constructed the following mixed-integer linear program (MILP) to find a flux vector, $v$ (with elements $v_r$ for each reaction $r$), log-transformed metabolite concentrations ($\ln(x)$) and binary variables indicating whether a reaction is feasible ($z$) given a specific objective function was satisfied. The objective function used in this work was to maximize biomass yield, similar to the objectives frequently used in FBA model of microbial metabolism. Thus, the optimization problem was constructed according the following MILP:

$$\begin{aligned}
\underset{\ln(x),v,z,e}{\text{maximize}} \quad & v_{\text{biomass}} \\
\text{subject to} \quad & Sv = 0 \\
& 0 < v_r \le z_r ub_r,\ \forall r \in \mathcal{R} \\
& z_r K - K + \Delta_r G' < 0,\ \forall r \in \mathcal{R} \\
& \Delta_r G'^o + RT\sum_i s_{ir}\ln(x_i) + \sigma_r e_r = \Delta_r G' \\
& \ln(10^{-6}) \le \ln(x_i) \le \ln(10^{-1}),\ \forall i \in \mathcal{M} \\
& -\sigma_m \le e_r \le \sigma_m\ \forall r \in \mathcal{R}
\end{aligned} \qquad [3]$$

where $\mathcal{R}$ and $\mathcal{M}$ are the sets of all reactions and metabolites, respectively. As discussed in detail elsewhere (13), the first equation in the constraint set ensures that intracellular metabolite concentrations are at steady-state, and are simply mass balance constraints for each metabolite. The second equation sets the bound on individual reaction fluxes, where the maximum flux through reaction $r$ is $ub_r$. Note that when $z_r = 0$, the flux through reaction $r$ is constrained to 0. The third equation sets ensures that $z_r = 1$ if and only if $\Delta_r G' < 0$, and $z_r = 0$ otherwise. Note that $K$ is a large number $(K > \max_r\{\Delta_r G'\})$ ensuring that this constraint is not violated with $z_r = 0$. The fourth equation is the free energy of each reaction as a function of log-metabolite concentrations. Note that we also add slack variables, $e_r$, to account for the possible error in the estimating standard molar reaction free energies for each reaction (where $\sigma_r$ is the standard error for each reaction $r$), which are bounded by a global error tolerance $\sigma_m = 2$ (set in equation 6). Lastly, equation 5 simply constrains the log-metabolite concentrations to be bounded between $1\mu$M and 100 mM. After each simulation, we performed a secondary optimization to find the minimal set of reactions that achieve the optimal growth rate by minimizing the $l_1$-norm of the flux distribution subject to the constraint that $v_{\text{biomass}} = v^*_{\text{biomass}}$

Numerical simulations were performed using the COBRA toolbox (16) and the Gurobi optimizer (Version 7.0.1). All source code is provided in the following github repository: http://www.github.com/jgoldford/protometabolic_modeling.

**Calculation of coenzyme and sequence-level features within enzymes.** To determine which reactions were associated with specific coenzymes (for results presented in Fig. S1B,D) we downloaded information for each Enzyme Commission number (E.C.) in the KEGG ENZYME database (http://www.genome.jp/kegg/annotation/enzyme.html). We downloaded each page and parsed the "comment" field for each E.C. and performed a text-based search to identify coenzymes associated with each E.C. number. We searched for text indicating that the enzyme mechanisms used one of the following coenzymes, cofactors and iron sulfur clusters: biotin, heme, PLP, TPP, pterin, molybdopterin, flavin, Fe, Co, Ni, Cu, Mn, W, Zn, Mo, Mg, FeS, FeFe, $Fe_2S_2$, $Fe_3S_4$ and $Fe_4S_4$, respectively. We also searched E.C. numbers indicating that the reaction mechanisms are non-enzymatic. Text-based searches were pruned manually to remove mis-annotated enzyme-coenzyme relationships.

For results presented in S1B, we computed the fraction of reaction E.C. numbers that were associated with one of the following coenzymes: Fe, Co, Ni, Cu, Mn, W, Zn, Mo, Mg, FeS, FeFe, $Fe_2S_2$, $Fe_3S_4$ and $Fe_4S_4$, or was marked as non-enzymatic. For results presented in S1D, we computed the fraction of reaction E.C. numbers that were associated with one of the following coenzymes: biotin, heme, PLP, TPP, pterin, molybdopterin, and flavin.

For results presented in Fig. S1E, we obtained a database of known enzyme active site residues (17). We first mapped the network reactions to E.C. numbers listed in KEGG, then identified active sites corresponding to E.C. numbers within the the expanded network. We next computed the fraction of active site residues containing nitrogenous side-chains, derived from the following amino acids: Arginine (R), Lysine (K), Glutamine (Q), Asparagine (N), Histidine (H), and Tryptophan (W).

### Supporting Information Text

**Network enzymes retain features of nitrogen-free catalysts.** In order for the expanded networks presented in the main text to have operated in prebiotic conditions, reactions would have been catalyzed non-enzymatically by inorganic or simple organic catalysts available in prebiotic environments. Prior work has suggested that reactions in metabolic networks that proceed spontaneously or depend on enzymes with inorganic coenzymes, such as iron-sulfur or transition metal cofactors, may have operated in prebiotic conditions (11, 18–20). We identified reactions in KEGG that could proceed spontaneously or are dependent on one of several inorganic coenzymes (Methods), and defined this set of reactions as *plausibly pre-enzymatic*, or "PPE"-reactions (Fig. S1A). For each proposed prebiotic scenario that lead to expansion of at-least 100 metabolites ($n = 144$, Fig S1A), we partitioned reactions added to the network before the inclusion of ammonia into the seed set (herein called "pre-ammonia" reactions) and reactions added to the network after ammonia was added to the seed set (or "post-ammonia" reactions). We then computed the fraction pre- and post-ammonia reactions that were classified as PPE reactions, and found that pre-ammonia reactions contained a higher proportion of PPE-reactions relative to post-ammonia reactions (one-tailed Wilcoxon sign-rank test: $P < 10^{-19}$), suggesting that the pre-ammonia reactions may have been more readily catalyzed by simple inorganic catalysts in prebiotic environments. We next hypothesized that if these enzymes evolved from a thioester-driven proto-metabolism without nitrogen, then enzymes in these networks should be depleted in enzyme-bound nitrogen-containing coenzymes. We thusly computed the fraction of pre- and post-ammonia reactions dependent on enzymes containing TPP, PLP, heme, biotin, flavin, pterin, and cobalamin (Fig. S1C, Methods). We found that the proportion of pre-ammonia reactions associated with these coenzymes were significantly less than the proportion of post-ammonia reactions dependent on these coenzymes (Fig. S1D, one-tailed Wilcoxon sign-rank test: $P < 10^{-24}$), which is primarily due to the large number of PLP-dependent reactions added to the network after the inclusion of ammonia.

Since only a minority of reactions in this network were categorized as PPE, simple organic or organosulfur catalysts may have been necessary in order for this network to function in prebiotic environments. Christian de Duve suggested that thioester-based polymers may have provided the necessary catalytic components of ancient metabolism in addition to inorganic catalysts (14). In modern living systems, monomers of keto acids are converted into amino acids, which are then polymerized into polypeptides either with or without the aid of the ribosome and mRNA. If prebiotic environments were severely nitrogen limited, keto acids may have been reduced to hydroxy acids, and polymerized into polyesters using thioesters as a condensing agent. Notably, in such a scenario only the polymer backbone is altered, leaving the side chains ($R$-groups) within today's amino acids intact. Recent work has demonstrated that polyesters may aid in the polymerization of amino acids during dry-wet cycles (21), and that the peptidyl-transferase domain on the ribosome can polymerize hydroxyacylated tRNAs to form polyesters (22, 23), suggesting that ester bond formation may have proceeded amide bond formation in living systems.

It has been proposed that enzymes retain features of early catalysts before the emergence of the genetic code and protein translation systems, and that enzyme active sites may bear resemblance to ancient catalysts. Thus, if this network represents a relic of an ancient metabolism before the biological incorporation of nitrogen, then the active sites of enzymes catalyzing reactions within the network should be depleted in amino acids with side chains containing nitrogen (Fig. S1E). To see if the catalytic residues of the enzymes in the pre-ammonia network were depleted in amino acids with nitrogenous side chains, we first obtained a database of catalytic site residues inferred from protein structures (17). After removing entries with interactions mediated by the peptide backbone, this dataset consisted of 18,721 entries, 1,304 of which were associated with active sites of enzymes in the nitrogen-free network in a representative network. For each putative prebiotic scenario resulting in an expansion with more than 100 metabolites, we computed the fraction of active site residues that contained nitrogen in enzymes associated with both pre- and post-ammonia reactions (Fig. S1E). We found that the proportion of nitrogenous catalytic residues associated with pre-ammonia reactions was significantly lower than the proportion of nitrogenous catalytic residues associated with post-ammonia reactions (Fig. S1F, Wilcoxon sign-rank test: $P < 10^{-24}$).

One potential alternative explanation for these biases in amino acid composition within the active sites of extant enzymes may be the outcome of evolutionary selection: nitrogen limitation in the environment may have favored mutations that lead to less nitrogen within these enzymes. However, evidence for selection for less nitrogen usage would manifest within the entire protein sequence, rather than just the active sites. Thus, we computed the fraction of amino acids with nitrogenous side chains across the entire coding sequences, rather than specifically the active sites, for enzymes associated with pre- and post-ammonia reactions (see Methods). We found no evidence that enzymes in the pre-ammonia network had a decreased usage of amino acids with nitrogenous side chains side chains relative to enzymes added to the network after ammonia was included in the seed set (one-tailed Wilcoxon sign rank test: $P = 1$) , suggesting that the biases within the active sites are not merely a consequence evolutionary selection (see Fig. S4).

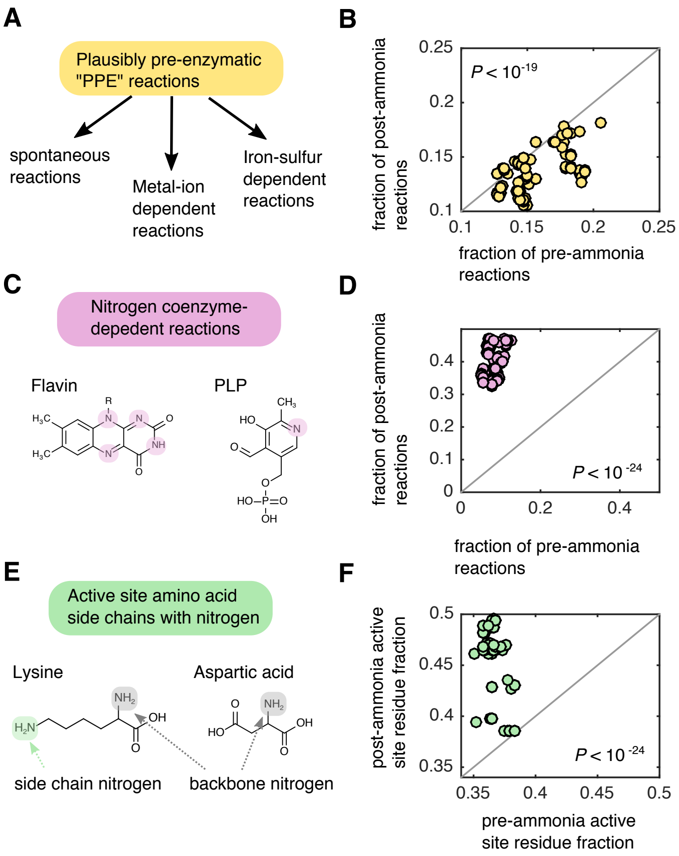


**Fig. S1. Enzymes in thioester-driven protometabolism are depleted in nitrogenous compounds** (A) We classified reactions in KEGG as being plausibly pre-enzymatic (PPE) reactions if they could (a) proceed spontaneously, (b) were associated with enzymes that contain at-least one iron-sulfur cluster or (c) were associated with an enzyme that relied on at-least one metal (Ni, Co, Cu, Mg, Mn, Mo, Zn, Fe, W) cofactor. (B) For all scenarios resulting in expansion of $> 100$ metabolites ($n = 144$, Fig. S1A) we computed the fraction of PPE-reactions amongst the pre-ammonia reactions (x axis) and post-ammonia reactions (y-axis). The frequency of PPE-reactions in the pre-ammonia reaction set was on average higher than the frequency of PPE-reactions in the post-ammonia reaction set (one-tailed Wilcoxon sign-rank test: $P < 10^{-19}$). (C) We identified KEGG reactions that were dependent on at-least one of the following nitrogen-containing coenzymes: flavin, biotin, thiamine pyrophosphate (TPP) pyridoxal phosphate (PLP), heme, pterin or cobalamin. (D) We compute the fraction of pre- and post- ammonia reactions associated with nitrogen containing coenzymes in the KEGG database, and found that a much higher proportion of post-ammonia reactions were dependent on these coenzymes relative to pre-ammonia reactions (one-tailed Wilcoxon sign-rank test: $P < 10^{-24}$). (E) We parsed the catalytic active site database (17) to find entries associated with pre and post-ammonia reactions, and compute the fraction of entries associated with amino acids with nitrogen-containing side chains (Q,N,W,H,K,R). (F) For each scenario, the fraction of active sites with nitrogen-containing amino acids was significantly higher for post-ammonia reactions relative to pre-ammonia reactions (one-tailed Wilcoxon sign-rank test: $P < 10^{-24}$).

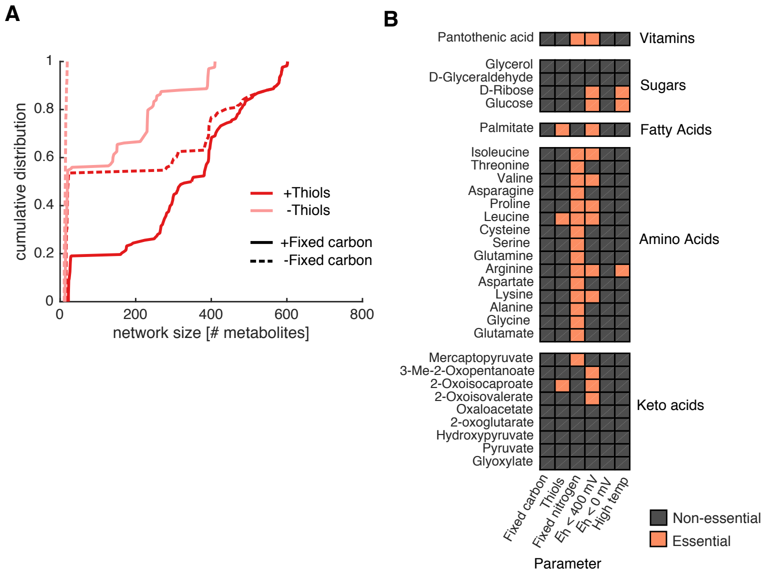


**Fig. S2. Thiols are required for autotrophic expansion and fatty acid production** (A) We grouped the $n = 672$ geoechemical scenarios into wither a source of fixed carbon and thiols was provided in the seed set. We then plotted the empirical cumulative distributions for each group of scenarios. Notably, when thiols and fixed carbon are not supplied in the seed set, the networks are always below 100 metabolites, indicating that expansion is prohibited without either fixed carbon or thiols in the seed set. (B) We determined what geochemical parameters ($x$-axis) were essential for the production of important biomolecules ($y$-axis). For example, palmitate, a long chain fatty acid, is producible only if thiols and reductant below 400 mV is provided in the seed set.

 

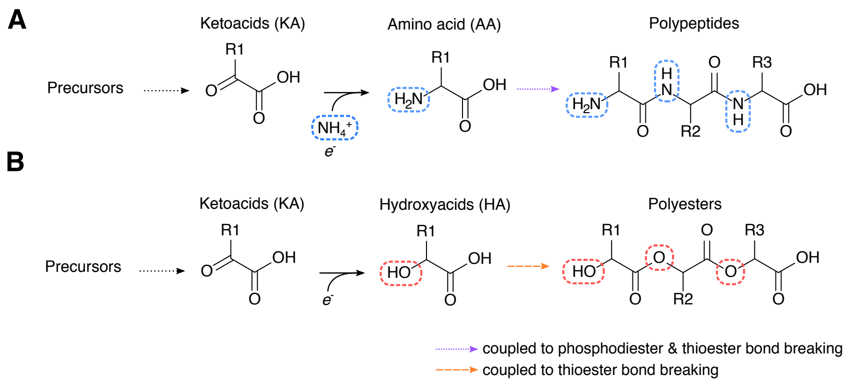


**Fig. S3. Putative ancient catalysts**. (A) In extant biochemistry, keto acid are converted to amino acids using transamination or reductive amination reaction mechanisms, which are then polymerized using a phosphate or thiester coupled mechanism to make polypeptides. (B) If prebiotic environments did not have a source of fixed nitrogen, then keto acids could have been reduced to $\alpha$-hydroxy acids, which could then be polymerized into polyesters either with (14) or without (21) thioester bond breaking.

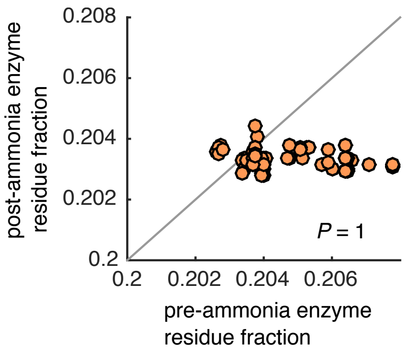


**Fig. S4. Enzymes catalyzing reactions before the addition of ammonia are not depleted in nitrogen containing amino acids relative to enzymes added after ammonia** To see if the amino acid biases in active sites of enzymes catalyzing reactions added to the network without ammonia (see Fig. S1E-F) is confounded due to evolutionary selection for reduced nitrogen in these enzymes, we computed the fraction of nitrogen side chains in enzymes in pre-ammonia reactions ($x-$axis) and in enzymes in post-ammonia reactions $y$-axis. We found that enzymes in the pre-ammonia networks did not have significantly less nitrogen usage compared to enzymes in post-ammonia reactions (one-tailed Wilxocon sign-rank test: $P = 1$).

**Additional data table S1 (scenario_results.csv)**

Results of the systematic exploration of 672 prebiotic scenarios. Columns labeled with KEGG compounds IDs denote whether a compound appeared after expansion (1) or not (0).
